# High-Throughput Stool Metaproteomics: Method and Application to Human Specimens

**DOI:** 10.1101/2020.03.06.981712

**Authors:** Carlos G. Gonzalez, Hannah C. Wastyk, Madeline Topf, Christopher D. Gardner, Justin L. Sonnenburg, Joshua E. Elias

## Abstract

Stool-based proteomics is capable of significantly augmenting our understanding of host-gut microbe interactions. However, in comparison to competing technologies such as metagenomics and 16S rRNA sequencing, it is under-utilized due to its low throughput and the negative impact sample contaminants can have on highly sensitive mass spectrometry equipment. Here, we present a new stool proteomic processing pipeline that addresses these shortcomings in a highly reproducible and quantitative manner. Using this method, 290 samples from a dietary intervention study were processed in approximately 1.5 weeks, largely done by a single researcher. These data indicated a subtle but distinct monotonic increase in the number of significantly altered proteins between study participants on fiber- or fermented food-enriched diets. Lastly, we were able to classify study participants based on their diet-altered proteomic profiles, and demonstrated that classification accuracies of up to 89% could be achieved by increasing the number of subjects considered. Taken together, this study represents the first high throughout proteomic method for processing stool samples in a technically reproducible manner, and has the potential to elevate stool-based proteomics as an essential tool for profiling host-gut microbiome interactions in a clinical setting.

**Importance:** Widely available technologies based on DNA sequencing have been used to describe the kinds of microbes that might correlate with health and disease. However, mechanistic insight might be best achieved through careful study of the dynamic proteins at the interface between the foods we eat, our microbes, and ourselves. Mass-spectrometry-based proteomics has the potential to revolutionize our understanding of this complex system but its application to clinical studies has been hampered by low-throughput and laborious experimentation pipelines. In response, we developed SHT-Pro, the first high-throughput pipeline designed to rapidly handle large stool sample sets. With it, a single researcher can process over one hundred stool samples per week for mass spectrometry analysis, roughly 10 times faster than previous methods. Since SHT-Pro is fairly simple to implement using commercially available reagents, it should be easily adaptable to large-scale clinical studies.

## Introduction

The gut microbiome is characterized by numerous complex interactions influencing human health and disease, and has been associated with disorders ranging from inflammatory bowel disease to autism^1,2^. Stool is a biologically rich biomaterial, containing host, microbe, and dietary proteins, among a rich array of biomolecules. The broad proteinaceous representation of relevant biological entities and interactions, in conjunction with non-invasive sample collection, makes stool ideal for studying the complex ecosystem at the host-gut microbe interface^3^. Microbiome composition can be readily determined using 16S rRNA gene sequencing from stool DNA, and due to its high-throughput nature, is well-suited for surveying a single individual over extended longitudinal time courses. Metagenomic and metatranscriptomic sequencing technologies can elucidate microbes’ functional capacities and states^4^. However, additional measurements are needed to elucidate the microbiome-host interactions that can profoundly affect host health.

Stool proteomics offers the ability to simultaneously measure both host- and microbe-expressed proteins, their post-translational modifications, and the dietary components also present in the gut. These components reflect interactions and physiological states that are otherwise difficult to survey through nucleic acid sequencing alone^5^. We previously showed that host proteins in stool reflect expression along the length of the gut and reveal signatures specific to type of inflammatory state, such as distinct levels of antimicrobial proteins. Importantly, these signatures can vary in a manner distinct from the gut microbiota^5^. For example, we showed previously that fecal microbiota transplanted into an antibiotic-induced *Clostridium difficile* infection mouse model normalized the microbial composition but not the host stool proteomic profile. Since proteins can be recovered from archived frozen stool samples, the approach offers a way to illuminate aspects of host mucosal biology non-invasively and longitudinally, long after stool collection.

Despite its functional utility, stool-based metaproteomics remains under-utilized compared to the aforementioned next generation sequencing technologies. One major hindrance to broader implementation has been low sample processing throughput. Indeed, while we and others previously demonstrated the power and utility of stool proteomics, those studies relied on data generated at rates as low as 10-30 samples per week^6–9^. Additionally, workflows developed for processing for cell culture and tissue lysates are not optimized to eliminate contaminating molecules that are abundant in stool. Insufficient contaminant removal can lead to instrumentation down-time, decreases overall sample throughput, and introduces experimental noise that can dilute biologically relevant signals.

Here, we describe a method, the Stool High Throughput Proteomics pipeline (SHT-Pro), that increases our ability to acquire high-quality metaproteomic stool analyses by as much as 100-fold when paired with multiplexing technologies such as tandem mass tag (TMT) labeling. As a first demonstration of this method, we applied SHT-Pro to 145 stool specimens longitudinally collected from 29 human participants as part of an ongoing dietary intervention study investigating the biological effects of diets enriched in fiber versus fermented foods (ClinicalTrials.gov Identifier: NCT03275662). Processing them in duplicate (290 total samples) using SHT-Pro took approximately one week from stool to mass-spectrometry-ready peptides, an estimated time savings of over 2.5 months compared to our previously published workflow^5,6^. The resulting data set identified over 5,600 unique host and microbial proteins, 45% which were shared between both study groups. We found that the number of proteins which significantly differed between the two groups increased over time, indicating that diet shapes the stool metaproteome of humans. We further demonstrate that the inclusion of more participants in metaproteomic analyses, in a fashion which this method enables, enhances the ability to classify study subjects compared to the smaller-scale datasets that were more feasible using prior methods. These data support SHT-Pro as overcoming a major hindrance for performing the kinds clinical-scale studies needed for statistically sound measurements of diet and its impact on the host and its gut microbiome.

## Results

### SHT-Pro increases sample processing speed with a high degree of reproducibility

A major limitation to large-scale adoption of stool metaproteomics has been is its heavily reduced throughput compared to DNA-sequencing. Considering the labor-intensive, multi-day nature of our previously published stool proteomics method, we found that one researcher could reasonably process 25 stool samples per week on average^6,10^. To narrow this gap, we developed our pipeline to rapidly process hundreds of stool samples (Fig. 1a) in a matter of days while maximizing liquid chromatography and mass spectrometry (LC-MS) instrumentation stability. 96-well protein traps columns (S-trap) for initial protein purification and digestion along with automation technologies for solid-phase extraction cleanup are two critical components of this added efficiency. To test time savings of the new method, we compared sample processing time of our previously published workflow to SHT-Pro (Fig. 1b). While processing fewer than 10 samples at a time does not result in substantial time savings (∼2.5-3 days saved), larger sample sets benefit from dramatic time savings. For example, processing 96 samples (a single 96-well plate) takes as little as 1.5 days using SHT-Pro, compared to approximately 30 days using the previous protocol (approximately 20-fold decrease).

**Figure 1.**
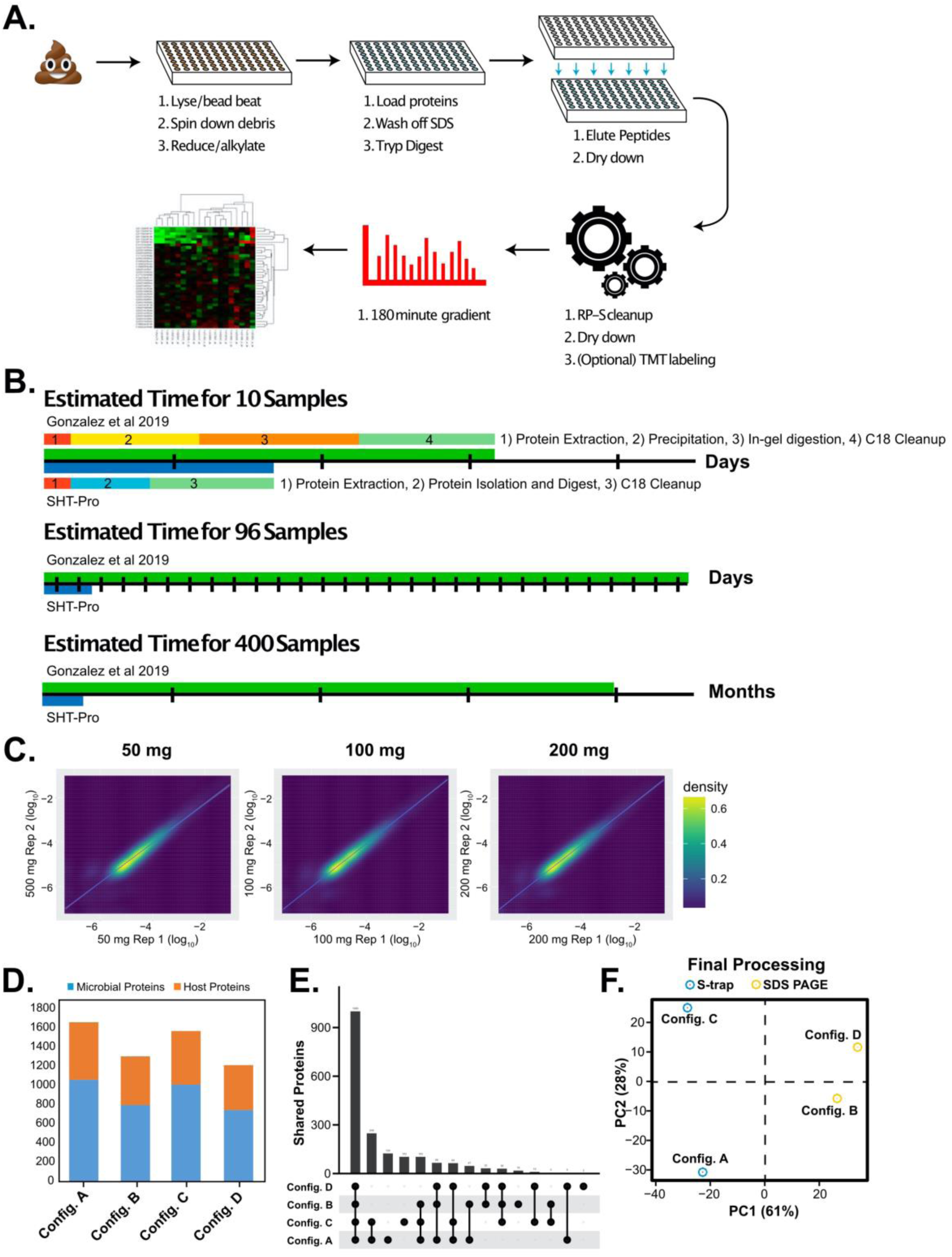
SHT-Pro is highly reproducible and reveals biologically relevant information. **A)** Simplified SHT-Pro processing pipeline. **B)** A comparison of estimated time taken to process varying amounts of samples. Multi-colored bars in the first section represent general pipeline stages. **C)** A scatterplot comparison of two replicate SHT-Pro analyses of a single stool specimen collected from an IBD patient during flare, using differing amounts of starting material but the same amount of sample loaded onto our chromatography columns (0.5 ug). **D)** Comparison of identified proteins from four different conditions: SHT-Pro (config. A), bead beating, TCA precipitation, and SDS-PAGE purification (config. B), vortex only, TCA precipitation, and SHT-Pro (config. C), or previous workflow (config. D). Each bar represents number of identified microbe or host proteins. See Table 1 for full experimental conditions. **E)** Shared subset plot of all four samples. **F)** PCA comparing the four SHT-Pro and previous workflow samples described in SI Figure 1C. Protein abundances were normalized and scaled (log_2_) prior to analysis.

Sample processing speed improvements would be of little value if the data quality of the resulting peptides were reduced due to poor contaminant removal. We found that repeated LC-MS analysis of a single stool specimen processed using SHT-Pro resulted in no degradation of LC-MS performance (often caused by sample contamination) as measured by analysis of the same standard peptide mixture interspersed throughout 20 SHT-Pro-prepared stool specimens (7,205 ± 60 unique peptides, n=4 replicates: one prior to stool LC-MS analysis, two spaced 10 stool analyses apart, one following 20 stool analyses) versus LC-MS analyses of the standard mixture on a new analytical column (average 7,350 ±150 unique peptides).

**Table 1.** Configuration of samples used in comparison of SHT-Pro and previous workflow. Table outlining the preparation conditions for each sample.

| Sample | Bead-beat | Vortex | TCA | S-trap | SDS-PAGE |
| --- | --- | --- | --- | --- | --- |
| Config. A | X |  |  | X |  |
| Config. B | X |  | X |  | X |
| Config. C |  | X | X | X |  |
| Config. D |  | X | X |  | X |

We next tested whether our preparation method also led to high reproducibility. To accomplish this, we aliquoted the stool specimen described above in varying amounts (50, 100, and 200 mg), and processed each aliquot by SHT-Pro in duplicate (SI. 1a). Starting material amounts were chosen based on our previous protocol as well as what we have found to be generally available from human clinical studies. We note, however, they do not test the lower limit of initial starting material needed for SHT-Pro. These analyses identified 11,373 unique peptides originating from 1,879 (1,152 microbial, 727 host) proteins, averaging 1,791 proteins per sample. Over 85% of all proteins were identified from all preparative replicates and starting material amounts, suggesting a low degree of sample preparation bias (SI 2a). All input protein amounts produced strong linear correlations (R^2^ values for 50 mg = 0.85, 100 mg = 0.92, 200 mg = 0.91; Fig. 1e). Similarly, comparing the intensity of proteins found in a replicate of the 50 mg samples to those of the 200 mg samples yielded an R^2^ value of 0.86 (SI 2b). As expected, these preparative replication correlation values were less than correlations between technical replicate LC-MS sampling from the same SHT-Pro-prepared peptide mixture (R^2^ = 0.99 ± 0.001, n=6 pairs). These data suggest a high degree of sample-to-sample processing fidelity.

We next examined how new procedural components of SHT-Pro compared to the lower-throughput aspects of our previously published workflow – specifically bead beating versus vortexing; and S-trap protein isolation versus TCA precipitation combined with SDS-PAGE. We evaluated the number of overall identifications made with two variations of each method, using parallel aliquots of the same stool sample described in Fig. 1a (see Table 1 for sample configurations). Combined, the four measurements yielded 2,352 total proteins identifications. Both samples processed with S-trap protein isolation and digestion had similar numbers of protein identifications to the SHT-Pro pilot detailed above (average of 1,610, ± 66). Samples processed with TCA precipitation and SDS-PAGE purification yielded less protein identifications (average of 1,255 ± 65). Of note, samples processed with the complete SHT-Pro yielded the greatest number of identifications (1,657; Fig. 1d). Comparing proteins found in this SHT-Pro sample to those from the initial pilot yielded an R^2^ value of 0.67 ± 0.014, despite originating from two different preparative replicates thawed and processed months apart.

The ratios of microbe-to-host protein identifications were slightly higher for SHT-Pro-prepared samples (mean = 1.8) than our previous workflow (mean = 1.5). Despite this larger proportion of microbe protein identifications, we found that the numbers of host proteins identified only with SHT-Pro (595, config. A) were substantially greater than the number of host proteins identified with our previous workflow (467, config. B). As with the experiment described in Figure SI 2a, the largest subset of proteins (1,000, 43% of all proteins) were present in all samples, regardless of preparation pipeline (Fig. 1e).The next largest unique protein set (248) included those shared only by the two S-Trap prepared samples (config. A and C) and not identified in the SDS-PAGE preparations (config. B and D). Only 32 proteins were found solely in SDS-PAGE-prepared samples (B and D). We attribute the decreased overlap (43% versus 85%) of this data set compared to the dilution-series sample set (SI 2a) to the differing sample preparation conditions of each experiment. The two samples that included SDS-PAGE were more alike in their proteomic profile when compared to the two samples that were processed with the S-Trap (A, C) (Fig. 1f). This is likely due to config. C’s use of TCA precipitation prior to S-trap processing, while config. A did not, which may cause an increase in specific protein subsets.

Given the improved speed, reproducibility, and sensitivity we observed with SHT-Pro, we next tested whether this method tended to identify proteins with biological relevance to the gut environment. We subjected the 100 most abundant proteins to gene ontology enrichment analysis using ShinyGO^11^. This revealed that the source stool specimen was significantly (FDR < 0.05) enriched for antimicrobial activity, neutrophil activation markers, as well as increased protease activity (SI. 3). Given the specimen set originated from a patient with inflammatory bowel disease during a flare, this result agrees with observations we^12^ and others^13^ have previously made, and supports SHT-Pro’s ability to produce biologically relevant information.

### Application of SHT-Pro to a longitudinal human diet study

The advances in throughput, coupled with high reproducibility and ability to reveal biologically relevant GI response pathways make SHT-Pro amenable to large sample sets which were previously impractical to process. To demonstrate SHT-Pro’s utility on a large, longitudinal data set, we applied it to samples collected from an ongoing dietary intervention study (ClinicalTrials.gov Identifier: NCT03275662). The overarching goal of this study is to elucidate how diets enriched in high fiber (e.g., whole grains, legumes, fruits) or in fermented (e.g., kombucha, kimchee, and yogurt) foods affect human health. Over the course of four months, study participants increased their intake of one of the two dietary intervention arms, and stool specimens were collected every two weeks over four phases: baseline, ramp-up (increasing intake), maintenance (peak intake), and choice (choose to eat respective diet, or not). We selected a subset of patients (n=29) for metaproteomic analysis based on sample availability during the baseline (two samples), ramp-up (single sample), and maintenance phases (two samples) for a total of five samples from each person over this period. The resulting 145 stool specimens were processed in duplicate, for a total number 290 stool measurements (Fig. 2a). Digested peptides resulting from initial processing with SHT-PRO were chemically labeled with tandem mass tag (TMT-11plex) multiplexing labels to increase throughput and quantifiability. Each TMT-11plex set contained one subject’s full time course in duplicate (5 timepoints * 2 replicates), plus one bridge channel representing a mixture of all 290 samples collected.

**Figure 2.**
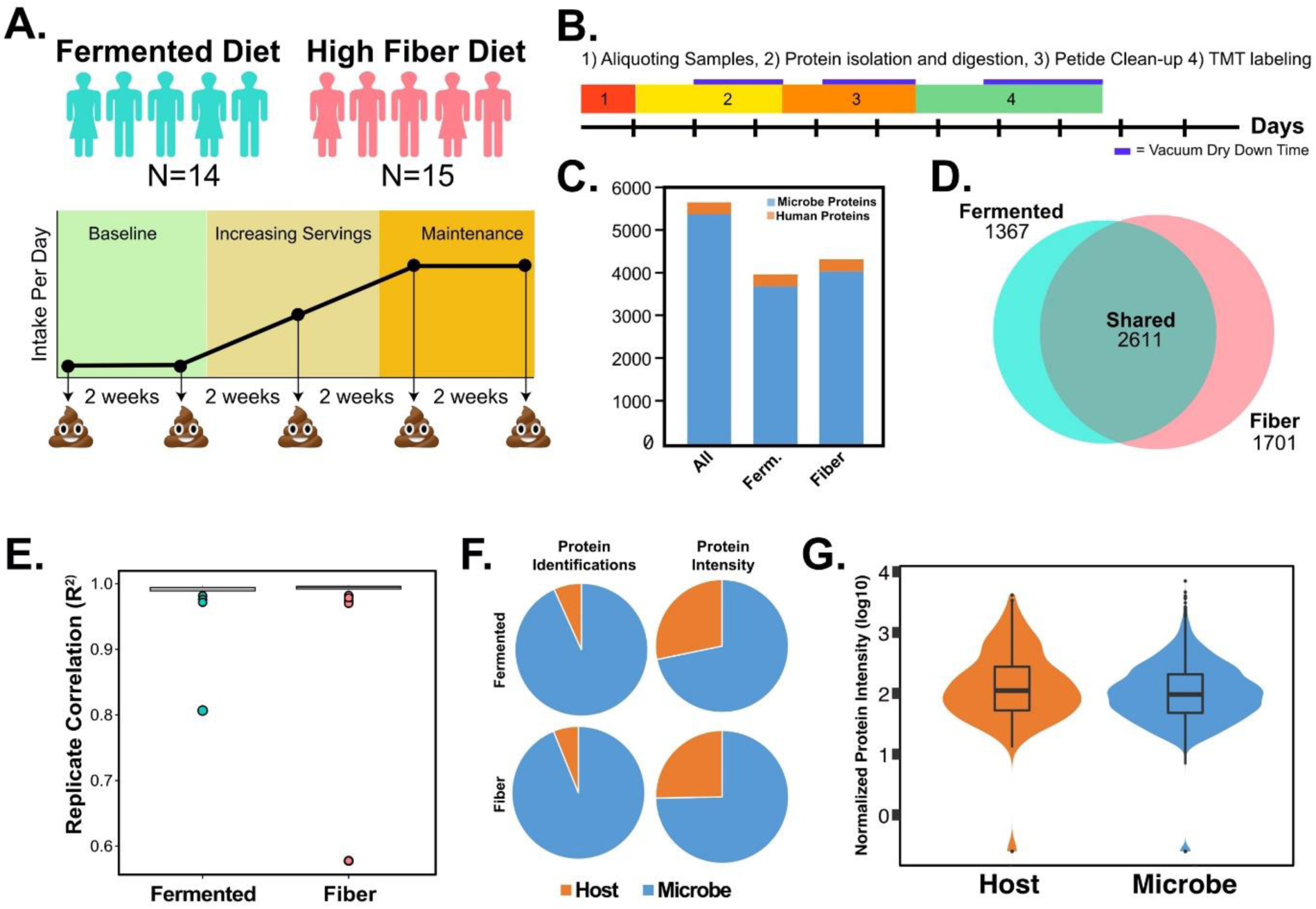
Diet study design and result overview. A) Illustration of major diet study components. B) Illustration of time taken for each step of SHT-Pro during the fiber and fermented group test. Section one and part of section two were done by three researchers, while 3 and four were done by a single researcher C) Bar graph of overall number of proteins identified as well as proteins identified within each diet group. **D)** Box plot of R^2^ values for fiber and fermented group (normalized and log2 transformed values). Average R^2^ value for fermented group was 0.997 while fiber group was 0.996. **E)** Venn diagram of shared and unique proteins identified in diet groups. F) Pie charts comparing protein identifications per group and organism (host or microbe) as well as percentage intensity of microbes and host proteins. G) Violin plot comparison of host protein intensity/microbial protein intensity using various scales. Blue bars use sum intensity for each protein (host or microbe) as input while the red bars use average intensity for each protein as input. The median and mean of those intensities are reported.

Once all 290 stool samples were transferred to four 96-well plates, they were then processed over the course of <9 days, including approximately 2.5 days devoted to TMT labeling and the associated cleanup steps that follow labelling. A single researcher carried out 80% of these steps (Fig. 2b). This sample set resulted in the identification of 83,061 high-confidence (q < 0.01) peptides (16,463 unique) assigned to 5,679 protein families (Fig. 2c; SI Tables 1, 2). Of these, approximately 94% (5,372) of identified proteins originated from microbes, with a much smaller host protein set (307). We found a large group of proteins (2,611, 46%) were shared by participants in both groups, while 54% (3062) of proteins identified were uniquely identified within just one dietary subgroup (fermented = 1,361, fiber = 1,701) (Fig. 2d). Replicate stool preparations were highly correlated (average R^2^ > 0.995 for both groups) with only two of 145 replicate pairs receiving an R^2^ value of < 0.90 (0.81 and 0.57), confirming a high degree of overall preparative reproducibility (Fig. 2e).

Having established the stability of SHT-Pro, we next focused on how microbial and host proteins compositionally contributed to the dataset at a high level. Despite comprising just 8% of all proteins identified, host-expressed proteins claimed much larger proportions of overall protein abundances (fermented = 25%, fiber = 28%) with an average host protein intensity approximately 67% (fermented = 72%, fiber = 63%) greater than their microbial counterparts (Fig. 2f-g). At the level of individual study participants, the fermented group had an average of 805 proteins in each sample, while the fiber group had 870 proteins (SI 5). This difference was not significant (p = 0.19, unpaired t-test), suggesting that both groups identified similar numbers of proteins despite the differences in diet. Together, these data demonstrate SHT-Pro workflow yields quantitatively consistent metaproteomic measurements when used with TMT labels or label-free quantification.

### SHT-Pro highlights presence of diet-responding proteomic subset

Having shown the efficacy of SHT-Pro in generating large and reproducible metaproteomic surveys, we next sought to understand if these two diets had any discernable effects on the stool proteome. Comparing microbe and host-expressed proteins via Principle Component Analysis (PCA, SI 6) suggested several global trends. First, we found that microbial protein variation across all study participants was largely explained by the first principle component (37%). Three participants within the fiber group were distinguished from the other participants by PC2. In contrast, host proteins exhibited less subject-specific clustering. Overall, neither microbial nor host protein measurements could clearly distinguish diet-induced effects at this high-dimensional level, whereas individual-specific microbial protein expression was much more substantial. This observation aligns with previous reports of microbiome composition profiles measured via 16S rRNA amplicon sequencing^14,15^.

Having observed minimal intergroup differences from high-level analysis, we next sought to determine whether biologically relevant temporal trends could be deduced at a more granular level. Comparing the proteomes of the two diet groups at each timepoint, we detected a trend suggesting diverging expression of both host and microbial proteomes subsequent to diet augmentation (SI 7b, Fig. 3a). More specifically, we observed an increase in significantly altered host (ramp = 9, maintenance = 17) and microbial (ramp = 10, maintenance = 45) proteins subsequent to the start of diet augmentations, although the significance of these small-number observations did not always a surpass strict (FDR < 0.05) multiple hypothesis testing(SI 7a-b). Nevertheless, a majority of these host proteins (14/17) increased among fermented group participants, while exhibiting negligible overall change in the fiber group. The STRING protein-protein interaction and gene-ontology platform suggested these 15 proteins were enriched (FDR < 0.05) in GO terms including maintenance of intestinal epithelium, GPI anchor binding, and sphingolipid metabolism (Fig. 3b). It is notable that nine of these 17 host proteins were also among the subset of 33 proteins identified in all study participants, regardless of diet group (Fig. 3d), and could therefore be useful markers of a wide range of host responses. As expected, the protein set common to all participants was strongly enriched (FDR < 0.05) in GO terms commonly found in the gut (Fig. 3c, SI table 4). These results suggest diet augmentation with fiber or fermented foods have a distinguishable impact on both host and microbial proteomes and highlights their ability to affect expression of highly prevalent gut-related proteins.

**Figure 3.**
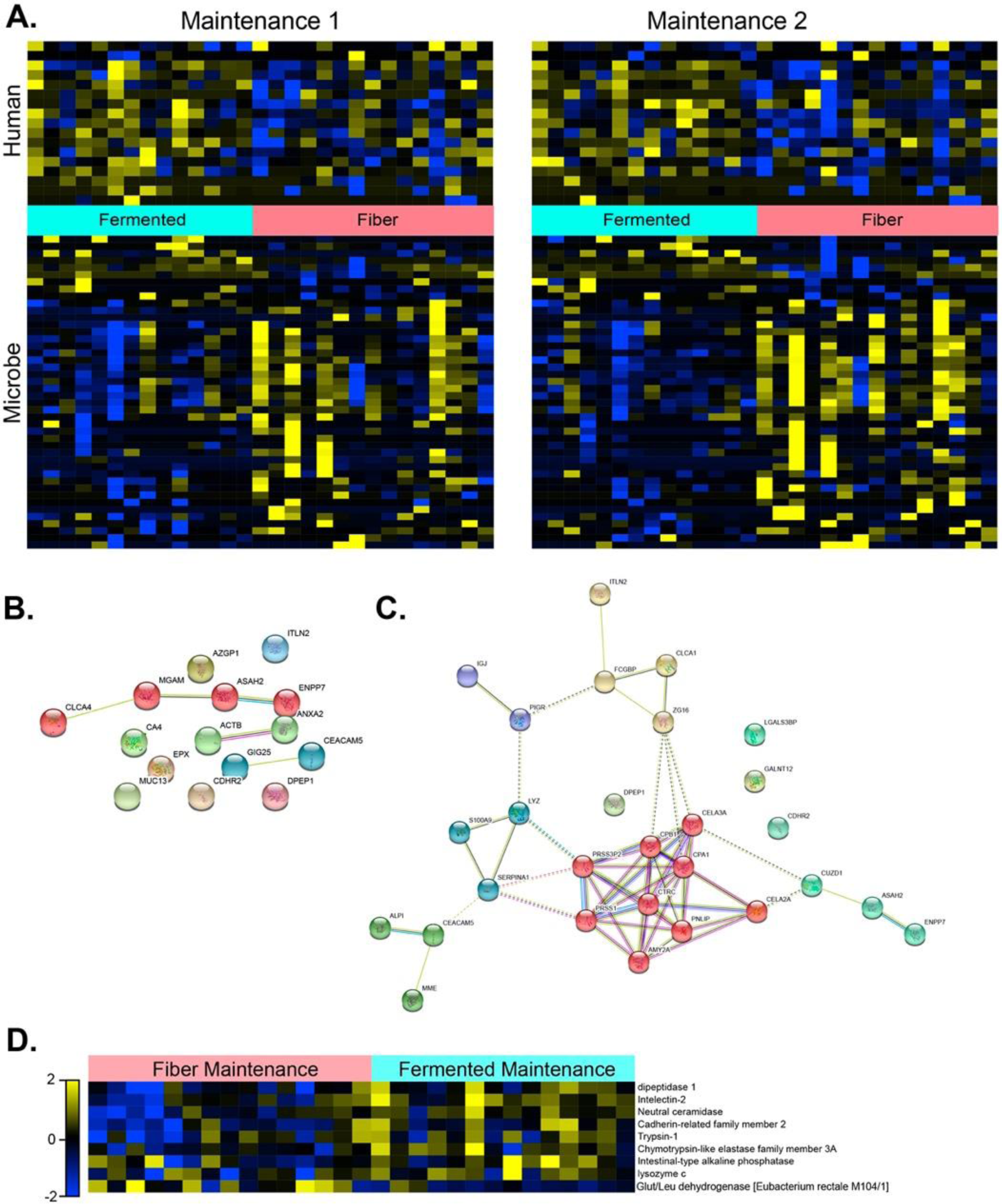
Diet intervention proteome reveals both unique and core proteome. **A)** Proteins that were significantly altered (p < 0.05) between fermented and fiber groups during the maintenance phase (timepoint 1 or 2). Proteins were also filtered for variation between participants (σ/σ_max variation_ > 0.15). See SI table 3 for accompanying list of significantly altered proteins and their normalized abundances. **B)** StringDB-generated functional network map using proteins significantly increased in the fermented group at the final maintenance timepoint. Nodes are colored according to the result of Markov-Clustering algorithm employed by StringDB, with each color signifying unique functional subnetworks. **C)** StringDB-generated functional network map of the commonly shared proteome. Colors overlaid on each node are the result of Markov-Clustering algorithm employed by StringDB. **D)** Universally identified proteins significantly (p < 0.05) altered during final day of maintenance period between fermented and fiber group diets. Proteins were normalized, log2 transformed, and scaled to levels present on the first baseline day (log_2day_ − log_2baseline_). Proteins were also filtered for variation between participants (σ/σ_max variation_ > 0.15).

These results indicate the stool proteome can be conceptually divided into several subset proteomes: an individual-specific proteome largely made of microbial proteins, a diet-impacted proteome, and a common ‘core’ proteome functionally associated with digestion and largely made up of host proteins common to most individuals. Given both common and unique proteome sets can be readily measured from all subjects, we next focused on whether these proteomes could be used to predict membership in either the fermented or fiber protein groups.

### SHT-Pro-derived protein abundance allow for classification based on diet group

As Figure 3 indicated, we observed modest diet-related differences between the fermented and fiber groups. However, we were curious as to whether more robust inter-group differences were obscured by considerable biological variation among this human cohort. To test this, we employed a leave-one-out cross-validated (LOOCV) random forest machine learning model, designed to identify distinguishing data features from complex, high-dimensional data^16^. The recursive feature selection approach we adopted chose differing combinations of study timepoints as model inputs, which were scaled to the first baseline timepoint (Fig. 4a).

**Figure 4.**
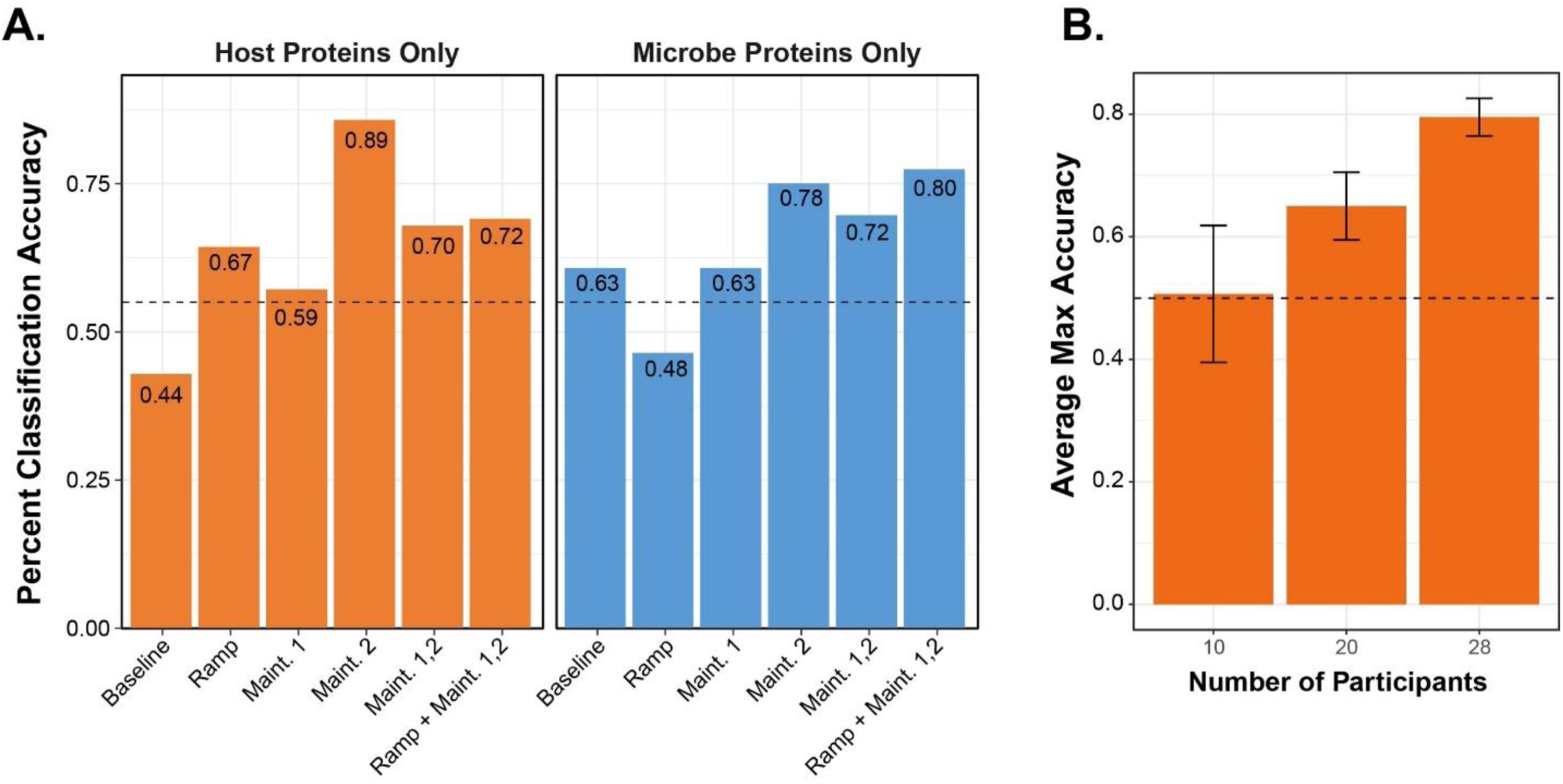
SHT-PRO-generated metaproteomes classify diet study participants. **A)** Bar plot highlighting classification accuracy of the random forest model using either microbial or host proteins from various combination of points along the time course depicted in Fig. 2a. Data were normalized by sample intensity and scaled to protein intensities found on the first baseline day. **B)** Bar plot comparing the LOOCV random forest model’s average maximum classification accuracy (e.g. average recorded maximum accuracy for each round of the ‘leave-one-out’ classification scheme) using only host proteins and varying the number of participants considered (n=10, 20, 28). Ramp and both maintenance phase days were used for each model while participant number was varied.

To gain insight into whether microbial or host protein abundance on specific days was of more predictive value in classifying these individuals based on their augmented diets, we ran the classifier only on host or on microbial proteins, considering either individual time points (e.g. only ramp, only maintenance day 1, etc.) or aggregated time points based on intervention status (e.g., all three post-diet intervention days) (Fig 4a). Overall, we found that the greatest classification accuracy was achieved by considering the abundance of solely host proteins from the final maintenance measurement (89%). In contrast, microbial proteins measured at this time point only yielded 78% accuracy. However, it is noteworthy that evaluating the ramp and both maintenance time points together improved this classification somewhat for microbial proteins (80%), but decreased classification accuracy for host proteins (72%). These data suggest a more comprehensive microbial profile measured following diet induction captured both transient and sustained diet-specific signals, whereas host protein expression tended to evolve over the course of the intervention.

Given that host proteins better predicted group membership (Fig. 4a), we next varied the number of participants included in the model (Fig. 4b) to test whether this observation depended on the underlying depth of the data set. As expected, we observed increased classification accuracy as more study participants were included in the model (averages of 51%, 65%, and 80% accuracy for 10, 20, and 28 participants, respectively) signifying the necessity of more extensive data sets for studies focused on disease prediction and explanatory power. Despite being less than 10% of each sample’s proteomic profile, these data support host proteins’ ability to generate a more accurate classifier than microbial proteins.

## Discussion

Stool-based proteomics’ potential use as a basic science tool and a rich resource for clinical biomarker discovery has been touted for over a decade^17–19^. However, its broad adoption has been largely hindered by multiple difficulties. Chief among them is processing large sample numbers in an efficient manner, a necessity for studies with large heterogenous populations such as human trials. While some new protein extraction and digestion protocols have robust sample processing pipelines, they targeted processing 5-25 samples per week and thus far lack the ability to scale via automation. This limits their usefulness to large, longitudinal clinical studies. Furthermore, several such protocols we have tried in our lab fail to remove major contaminating molecules found in stool, which is evident from the continuous fouling of liquid chromatography columns and unstable mass spectrometer performance^7^. Both necessitate increased equipment maintenance, and associated down-time^7^. SHT-Pro resolves these deficiencies by leveraging a workflow specifically designed for large longitudinal stool collections, while maintaining the flexibility to accommodate smaller sample numbers for pilot studies, all with a high degree of experimental reproducibility. Our previously published methods required the processing large sample sets over multiple months, leading to a greater need to distinguish prominent preparation artifact from desired biological protein profiles. Here, we show that SHT-Pro can produce highly reproducible data sets spanning hundreds of samples in a matter of days. For example, in the current study SHT-Pro saved an estimated 3.5 months (approximately 80% less time) compared to our previous protocol. Since it is compatible with multiplexing technology, LC-MS data generation times can be further accelerated by an additional order of magnitude. Similar to the high-throughput DNA sequencing pipelines used to characterized gut microbial communities on a massive scale, we envision SHT-Pro will modernize the stool proteomics field and allow the profiling of a variety of diseased conditions ranging from IBD to multiple sclerosis^20^.

Nevertheless, we acknowledge that SHT-Pro as described here could be further improved in several simple ways. First, the aliquoting of initial stool samples is presently the most labor intensive and time-consuming step of the overall process. In our diet study, each sample was aliquoted by hand from the original specimen collection vessel to the 96-well bead beating plate (Fig. 1a). Given that this is a common obstacle for DNA and protein sample preparation pipelines alike, the microbiome field would benefit from an aliquoting technology targeted at this sample handling burden. Next, while multichannel pipettes currently used in SHT-Pro were critical components, a 96-well pipettor (single head or as part of an automated system) could more uniformly and rapidly dispense buffers, thus increasing overall speed of the assay while decreasing the amount of hands-on time laboratory researchers must invest in an experiment, and decreasing preparative variation.. Lastly, we noted a large portion of time within SHT-Pro was spent evaporating and concentrating samples via Cetrivap/Speedvac vacuum-based concentrators (Fig 2b). Given the larger volumes 96-well plates produce using our method (100-300 uL/well), this can be a significant hindrance to overall throughput. As such, an alternative method to concentrate peptides would significantly increase the throughput of SHT-Pro, potentially bringing sample preparation time to less than a day.

In the current study, we observed over 5,600 proteins that that could hold new biological insights into the impact of dietary fiber versus fermented foods. While this identification depth is greater than some of the first reported metaproteomic searches on stool, newer studies have reported substantially more host and microbial protein identifications (53,000 total proteins, various biological matrices)^9,17,21^. We attribute the decreased number of identifications in our current study to several factors. First, our use of the TMT multiplexing reagent creates a bias towards proteins that are found in multiple samples: signals found just one sample are diluted by the number of channels used^22^. Thus, we suspect many low-abundant, sample-specific host and microbial proteins were not identified. To combat this, future iterations of SHT-Pro could incorporate peptide fractionation and longer mass spectrometry runs per sample, which has been shown to significantly increase identification of low-abundant host and microbial proteins^9^. In this context, striking a balance between throughput and proteomic depth is crucial, as the biological and health-related significance of low-abundance proteins remains promising but unclear. Next, while the database used to search these samples (adapted from the Human Microbiome Project) is fairly extensive, the use of subject-specific metagenomes for the generation of protein databases would likely increase sample- and subject-specific protein identifications. Lastly, in comparison to previously published work, we injected approximately 4x less material into the mass spectrometer (0.5 ug vs 2 ug)^21^. Given that on average, we collected approximately 60 ug of peptide from each sample (over 100 ug/sample was collected in the pilot study), injecting more peptide or fractionating samples would likely increase our protein identification rate.

Despite these remaining challenges, SHT-Pro-generated metaproteome data which resulted in biologically meaningful insights, even in the context of a largely uncontrolled human diet study. Indeed, SHT-Pro revealed a subtle divergence in proteomes after the introduction of fiber and fermented diets, as evidenced by the increased number of significantly altered host and microbial proteins during the ramp and maintenance phases, while baseline measurements remained largely unchanged. These significantly increased proteins were enriched for several categories including intestinal epithelium maintenance and host sphingolipid metabolism. Interestingly, sphingolipids along with chemical variants (e.g. glycosphingolipids) and derivatives, have been previously shown to regulate invariant natural killer T-cells (iNKT)^23^. More recently, *Bacteroides fragilis*, a common gut-dwelling microbe, has been shown to produce sphingolipids, and their production protected mice from a oxazolone-induced colitis model, an effect largely mediated by their regulation of iNKT activation^24^. Here, the introduction of fermented foods may have increase the levels of *B. fragilis*, as has been previously noted in rats fed fermented tempeh, which in turn may increase levels of sphingolipid availability^25^. In the current study, we observed 20 proteins attributed to *B. fragilis*, however they showed no significant abundance differences between diet cohorts on the final day of maintenance. It is possible that the search algorithm used (TurboSequest, Proteome Discoverer 2.2) was not ideally suited to attribute peptides (and proteins) to the correct species in such a large search space, a common problem in the metaproteomic field^3^. In this case, metaproteomic-centric search suites such as MetaLab may be of some benefit^26^. While the purpose of this manuscript is to showcase SHT-Pro as an integral facet necessary for understanding host-microbe interactions, this result suggests that future studies using SHT-Pro may also benefit from a multiomic approach that also leverages 16S rRNA amplicon sequencing and metabolomics profiling. Nevertheless, SHT-Pro-generated data is compelling when considering that, other than dictating increased intake of each experimental cohort’s respective diet, study participants had no other nutritional restrictions. As such, any changes in the microbial or host stool proteome could be expected to be subtle and subject specific, and likely hidden by data-driven noise. This subtlety is highlighted by the classification success, which was only possible using machine learning techniques and not easily discernable by simply focusing on simple abundance changes. Importantly, the observed success of the LOOCV random forest model also suggests future microbial proteomic studies would likely benefit from normalization to a participants’ unique baseline signature as well as the inclusion of many participants, an inherent strength of SHT-Pro.

These data likely harbor many more insights, including revealing components of diet. While we have not mapped dietary peptides in this study due to database limitations, plant peptides are evident in our dataset and suggest utility in helping inform the many challenging aspects of dietary assessment in free-living humans. When paired with other ‘omic’ data (e.g. 16S rRNA, metabolomics, clinical measurements) these proteomic profiles are poised to significantly contribute to our understanding of the dietary impact on individuals over time. However, it must be noted that the focus of this article is largely the increase in quality and speed of SHT-Pro compared to the previous workflow and more in-depth analysis of multi-omic data associated with the dietary intervention will be completed as part of a larger publication.

Nevertheless, taken together, SHT-Pro reveals itself as a robust pipeline for processing stool samples in an extremely timely manner, and we believe its widescale adoption and improvement will enable powerful discoveries in the field of host-gut microbiome interactions.

## SI Figs

**SI 1.**
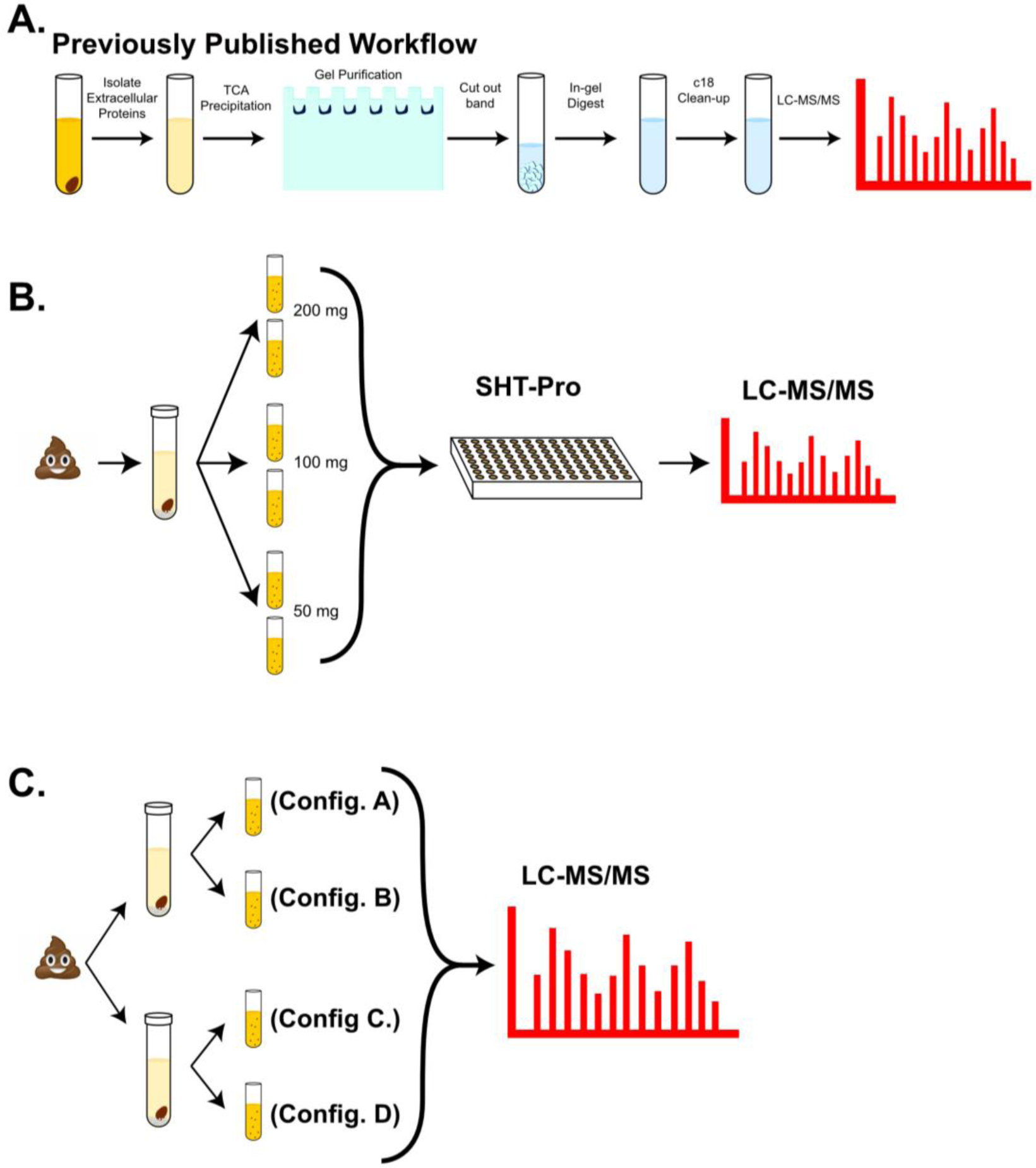
SHT-Pro experimental illustrations. **A)** Generalized experimental pipeline of our previous workflow. A more extensive description of the method is found in Gonzalez et al^6^. **B)** Experimental outline designed to test SHT-Pro reproducibility. More extensive experimental details are available in the methods section. **C)** Experimental outline designed to allow the comparison of the previous workflow to SHT-Pro. More extensive experimental details are available in the methods section.

**SI 2.**
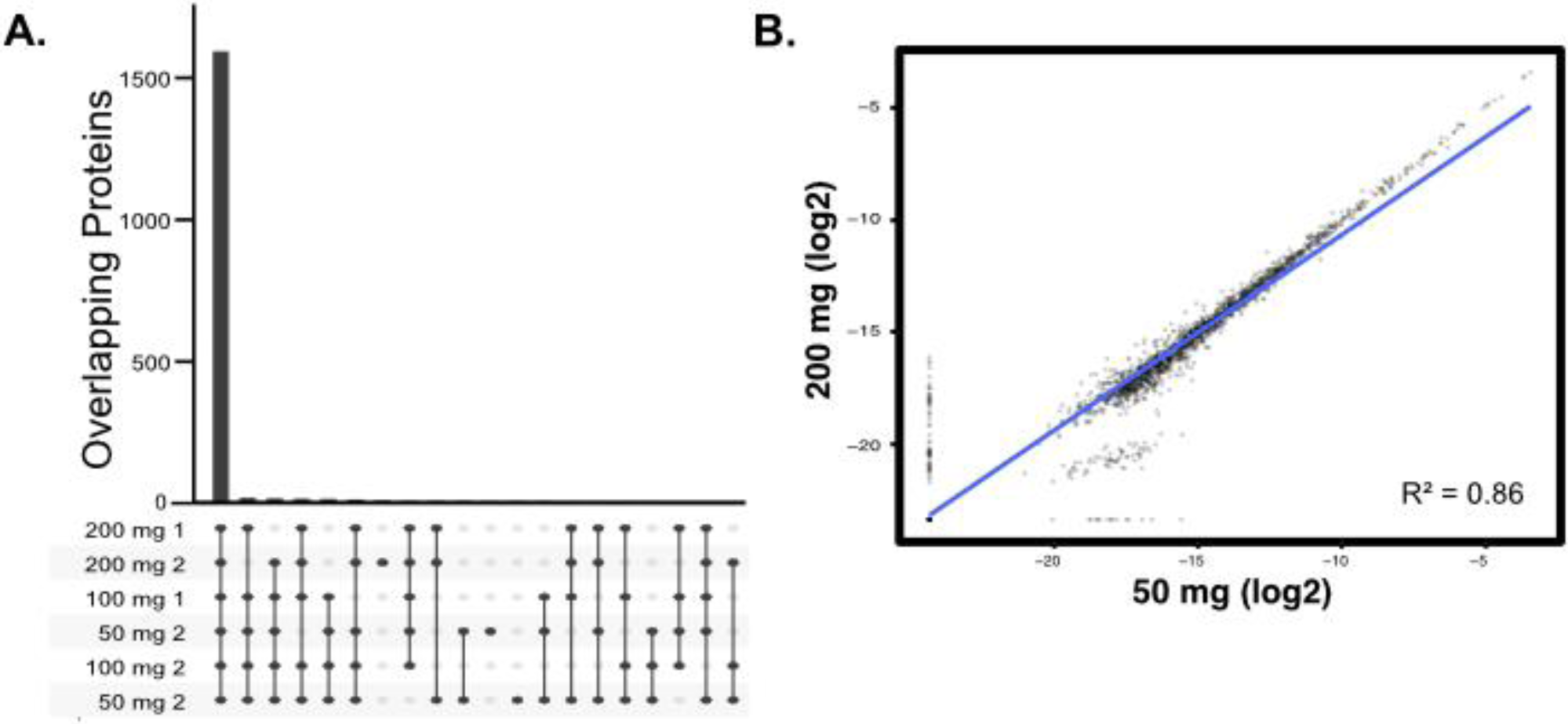
Pilot study protein comparisons. **A)** Plot generated using binary protein counts from each sample in the pilot experiment and the R package UpSetR. Dots connected at bottom of graph represent sets containing the number of proteins in the corresponding bar above the dots. **B)** Scatter plot comparison of a 50 mg (starting material) replicate vs. 200 mg (starting material) replicate.

**SI 3.**
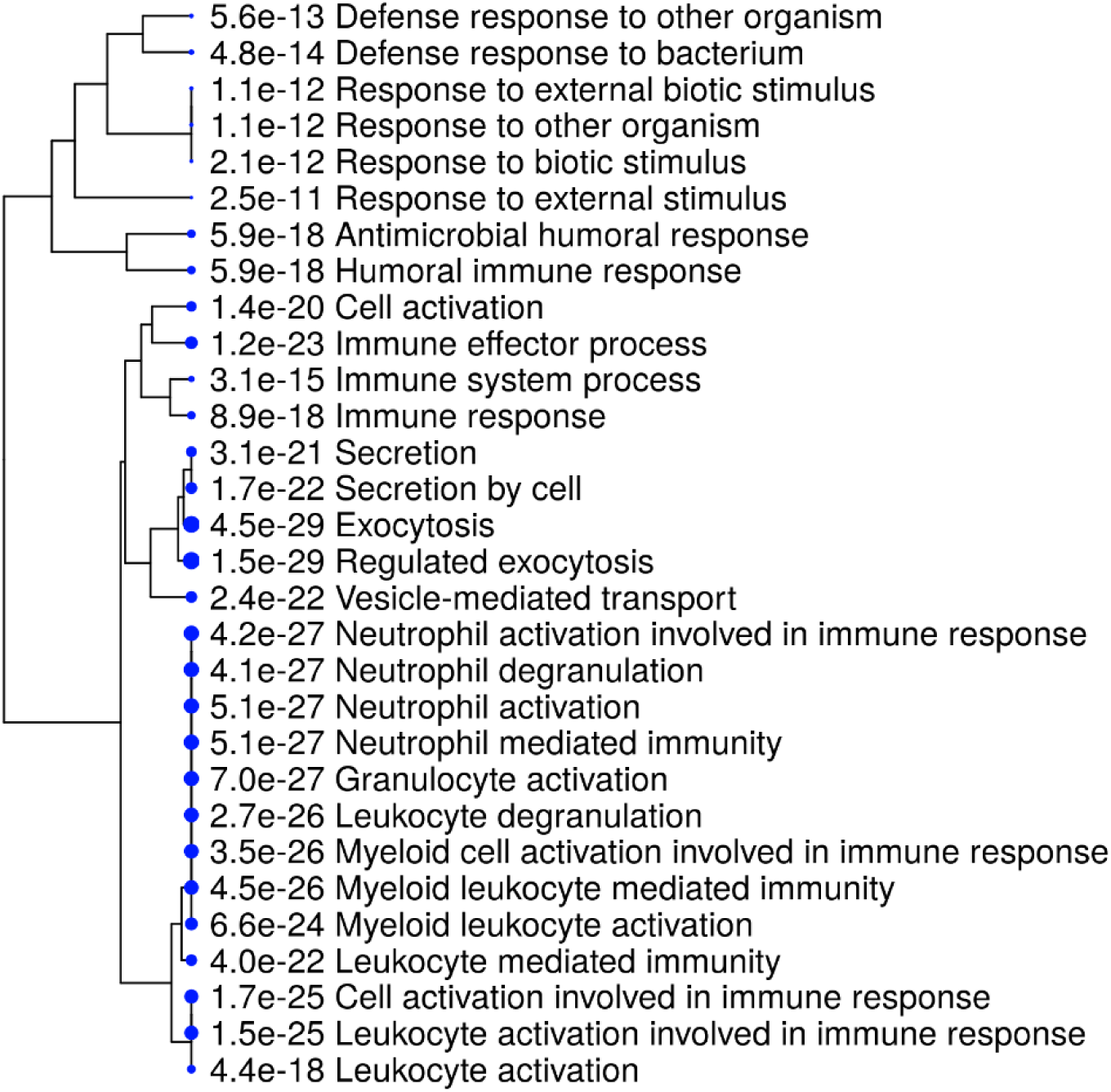
ShinyGo enrichment map. The top 100 overall most intense proteins were submitted to the gene ontology (GO) tool ShinyGO and clustered by biological similarity. Size of blue circle by GO term is proportional to its statistical enrichment score (FDR).

**SI 4.**
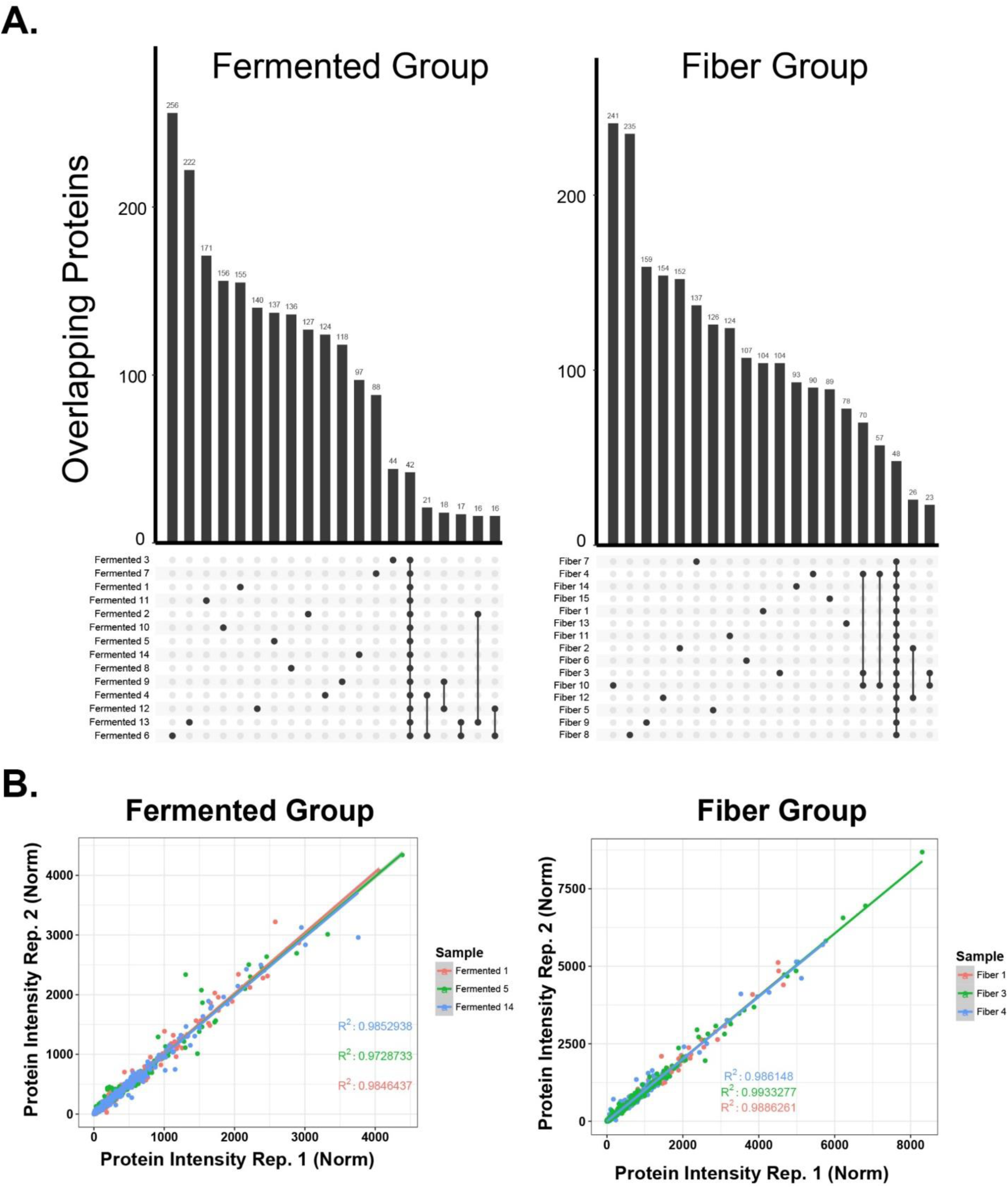
Protein sets by group and scatter plots of randomly chose fermented or fiber data sets. **A)** Plot generated using protein counts from each sample in the diet study and the R package UpSetR. Dots connected at bottom of graph represent samples corresponding to the containing protein count bar above it. **B)** Scatter plots of randomly chosen samples from SHT-Pro-generated data sets (3 fermented, 3 fiber). Samples are compared using normalized intensity and arbitrary units generated by Proteome Discoverer (v2.2) normalization feature.

**SI 5.**
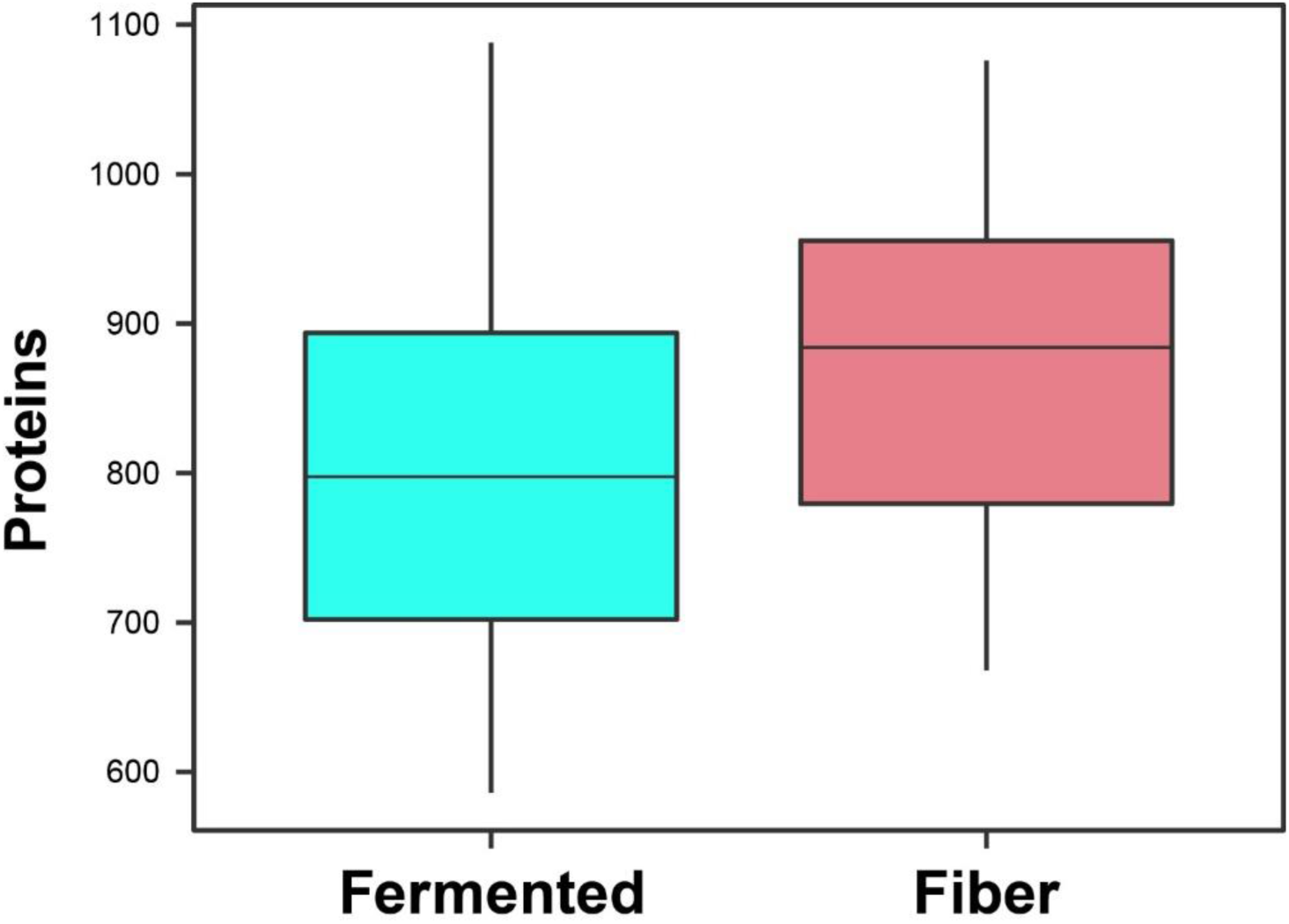
Proteins per individuals within each group. Boxplot of proteins identified in each individual broken out by diet group. No significant differences were found between the number of identified proteins in these two groups.

**SI 6.**
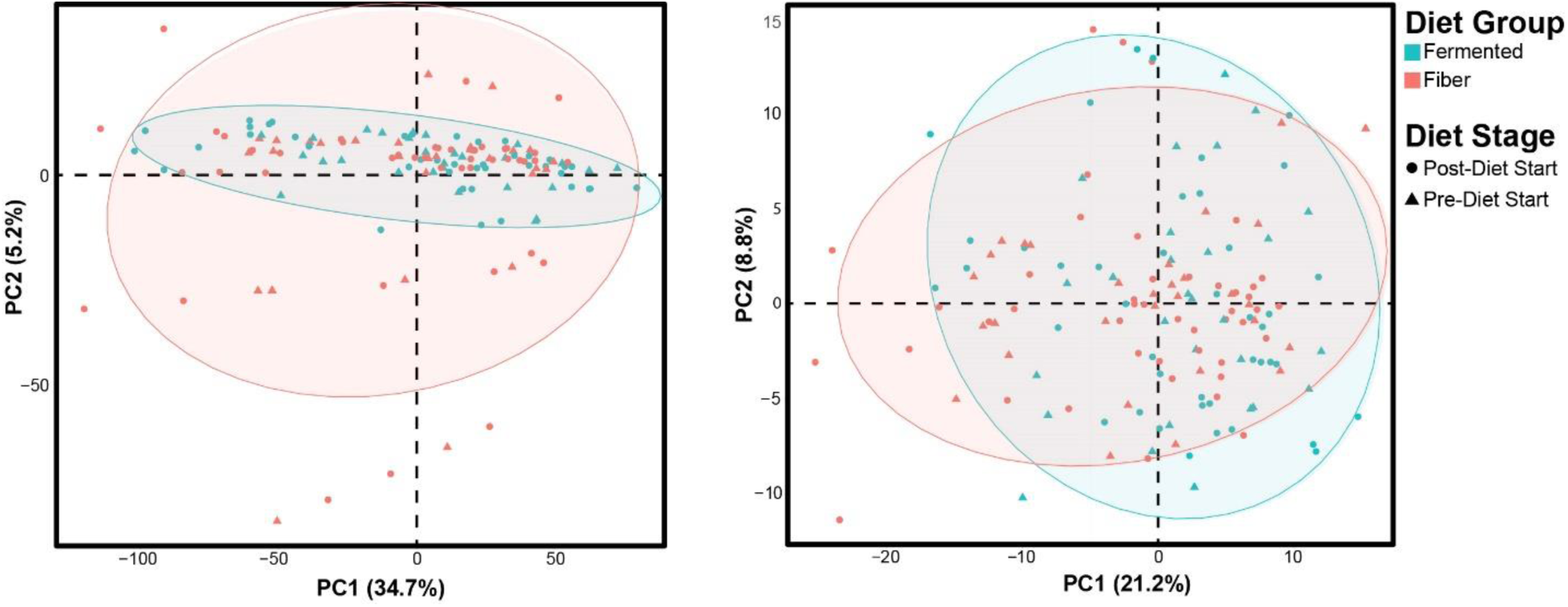
Proteins per individuals within each group. PCA of all microbial (5,372; left) or host proteins (307; right). Data used to generate these plots were normalized and log_2_ transformed. Ovals were automatically drawn to capture >85 % of data points within each diet study group.

**SI 7.**
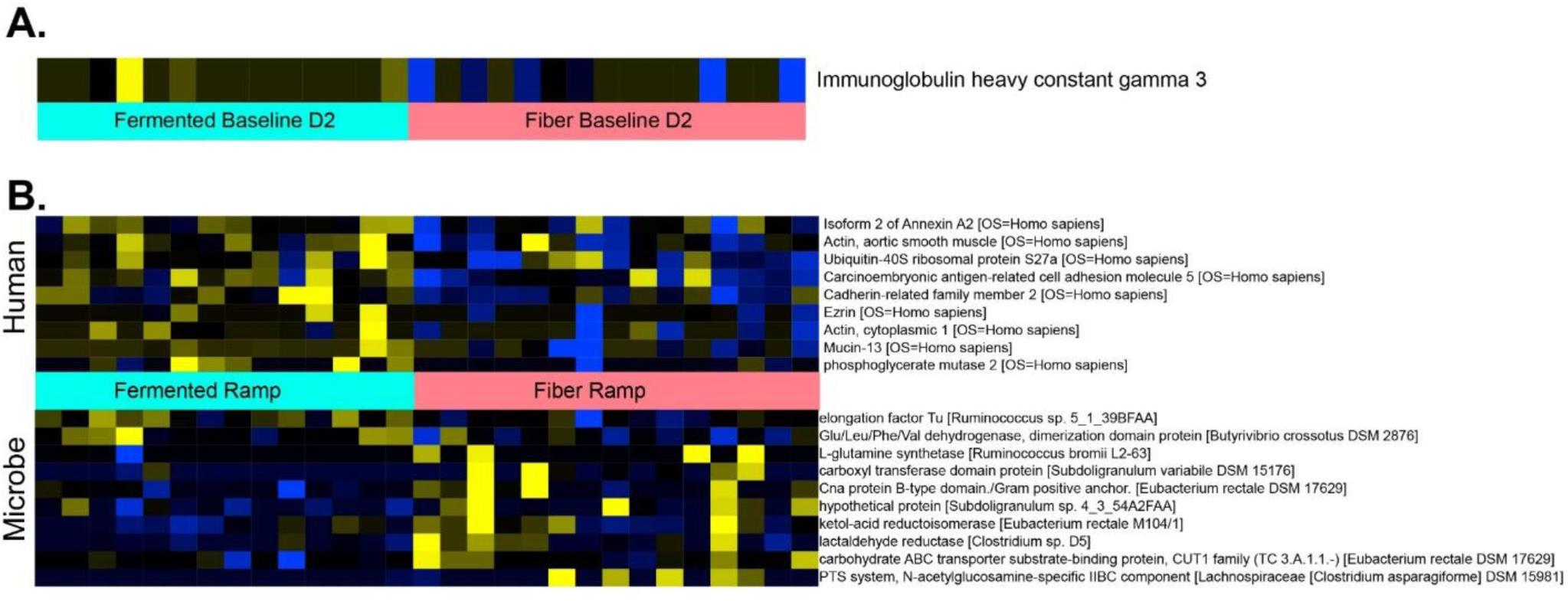
Heatmap comparison of significantly altered proteins within each stage of the diet study. **A)** One protein, an immunoglobulin heavy chain constant region, was significantly altered (p < 0.05) between fermented and fiber groups at baseline. No microbial proteins which significantly diverged between the two dietary cohorts were found at baseline. **B)** Proteins that were significantly altered (p < 0.05) between fermented and fiber groups during ramp phase.

## SI Tables

SI Table 1-Identified Peptides

SI Table 2 – Identified Proteins

SI Table 3 – Significantly altered proteins each day

SI Table 4 – GO term enrichment terms of core proteome (SI_core_bio_process.csv)

## Methods

### Buffers

#### Lysis buffer

6M urea, 5% sodium dodecyl sulfate (SDS), and 50 mM Tris were combined, with the pH adjusted to 8 using phosphoric acid. Roche cOmplete Mini protease inhibitor cocktail (04693159001 ROCHE) was added prior to adding buffer to samples.

#### Protifi Binding buffer (PBB)

90% methanol and 10% Triethylammonium bicarbonate buffer (TEAB, Sigma-Aldrich, catalog number T7408), adjusted to pH 7.1 using phosphoric acid.

#### Digestion buffer

100 mM TEAB and 5 ug trypsin (Promega V5113)

#### Peptide elution buffers

The first elution was performed using digestion buffer, the second elution was performed using 0.2% formic acid (FA), and the third elution was performed using 50% acetonitrile and 0.2% FA.

### Isolation of stool proteins and peptides (96-well variant)

Approximately 100-200 mg (when available) from each collected stool specimens was aliquoted into a 96 well plate along with approximately 600 mg of 0.1 mm ceramic beads (Omni International 27-6006). To each filled well, 750 uL of lysis buffer was added and plates were sealed with the OMNI-provided sealing mats. In order to increase their seal, each plate was additionally sealed with parafilm, although we found this was not necessary. The sealed plates were subjected to 10 minutes of bead beating at 20 Hz using an (Qiagen Tissuelyser II). After bead beating, each plate was centrifuged at 300 RCF at 4°C for 10 minutes. 500 uL of the resulting supernatant was transferred to a new 2 mL 96-well plate (Waters 186002482), sealed with a sealing mat and spun again at 300 RCF at 4°C for 10 minutes, then transferred into a fresh 2 mL plate. Samples were then reduced with 10 uL of 50 mM dithiothreitol (Sigma-Aldrich) for 30 minutes at 47°C, and alkylated with 30 uL of 50 mM Iodoacetamide (Sigma-Aldrich) for 1 hour at room temperature in the dark. 50 uL of the reduced and alkylated supernatant was transferred to a new 2 mL 96 well plate for further processing while the remaining material was stored at −80 for potential future analysis. Supernatant-resident stool proteins were washed, digested and eluted as described in the Protifi S-trap protocol (See http://www.protifi.com/wp-content/uploads/2018/08/S-Trap-96-well-plate-long-1.4.pdf for complete protocol). Briefly, the 50 uL of supernatant was acidified with 5 uL of 12% phosphoric acid to which 300 uL of S-trap binding buffer was added. Each resulting mixture was loaded into a single well. Positive pressure was used to load the proteins into each well (Waters Positive Pressure-96 Processor) with pressure at approximately 6-9 PSI on “Low-Flow” setting. **<u>Note:</u>** if after 1 minute, volume still remains in well, take a pipette tip and move any debris to side of well and it will begin to flow again. Loaded proteins were washed with 300 uL PBB five times. After washing, 125 uL of digestion buffer was added and proteins were digested for three hours at 47°C. Peptides were then eluted with 100 uL TEAB, followed by 100 uL of 0.2% formic acid, followed by 100 uL of 50% acetonitrile, 0.2% formic acid. These were captured in a 1 mL 96-well plate (Thermo Scientific AB-1127) and the volume was dried down in a Centrivap speedvac (Model 7810016). Plated samples were then desalted using RP-S tips on the Agilent Bravo AssayMap using built-in desalting protocol and eluted with 50% ACN and dried down. Plated peptide concentration was normalized using readings from the Biotek Synergy microplate reader and the Take3 microvolume plate, (single samples were adjusted using a nanodrop ND-1000). Samples were then labeled with TMT-11 multiplexing kit using the manufacture’s recommended method (Thermo-Fisher Scientific A34808). Channel-specific isobaric tag intensities were adjusted to 11(1:1) using recorded intensities from a 1-hour gradient mass spectrometry run and subsequently reinjected into the mass spectrometer after normalization.

### Isolation of stool proteins and peptides (individual tube variant)

Approximately 100-200 mg (when available) from each collected stool specimens was aliquoted into a bead beating tube along with approximately 600 mg of 0.1 mm ceramic beads (Omni International 19-732). To each tube, 750 uL of lysis buffer was added. Samples were subjected to 10 minutes of bead beating at 3500 RPM (OMNI Beadruptor 12 19-050). After bead beating, each sample was centrifuged at 300 RCF at 4°C for 10 minutes. 500 uL of the resulting supernatant was transferred to a fresh 2 mL tube, spun again at 300 RCF at 4°C for 10 minutes, then again transferred to a fresh 2 mL plate. Samples were then reduced with 10 uL of 50 mM dithiothreitol (Sigma-Aldrich) for 30 minutes at 47°C and alkylated with 30 uL of 50 mM Iodoacetamide (Sigma-Aldrich) for 1 hour at room temperature in the dark. 50 uL of the reduced and alkylated supernatant was transferred to a new 2 mL tube for further processing while the remaining material was stored at −80 for potential future analysis. Supernatant-resident stool proteins were washed, digested and eluted as described in the Protifi S-trap protocol (See http://www.protifi.com/wp-content/uploads/2018/08/S-Trap-mini-protocol-long.3.6.pdf for complete protocol). Briefly, the 50 uL of supernatant was acidified with 5 uL of 12% phosphoric acid to which 300 uL of S-trap binding buffer was added. Each resulting mixture was loaded into a single well. Vacuum manifold was used to load samples with pressure set at approximately 3-5 PSI. **<u>Note:</u>** if after 1 minute, volume still remains in well, take a pipette tip and move any debris to side of well and it will begin to flow again. Loaded proteins were washed with 300 uL PBB five times. After washing, 125 uL of digestion buffer was added and proteins were digested for three hours at 47°C. Peptides were then eluted with 100 uL TEAB, followed by 100 uL of 0.2% formic acid, followed by 100 uL of 50% acetonitrile, 0.2% FA. Eluate was captured and the volume was dried down in a Centrivap speedvac (Model 7810016). Dried down samples were then resuspended 250 uL 0.2% FA. Resuspended samples were then desalted using Seppak tC18 cartridges and subsequently dried down (Waters WAT036820). Each sample was then resuspended in 30 uL and the peptide concentration was normalized (Nanodrop ND-1000).

### Previous workflow protocol

Samples were prepared as described in Gonzalez et al^6^. Briefly, sample pellets were disrupted using 500 uL 8M urea lysis buffer supplemented with Roche cOmplete protease inhibitor (04693159001 ROCHE) by vortexing. After pellet resuspension, insoluble material was pelleted down at 2500 RCF for 10 minutes at 4C and the collected supernatant was subjected to ultracentrifugation (35000 RPM for 30 min. at 4C, Beckman-Coulter Optima Ultracentrifuge) to remove bacteria. The ultracentrifuge supernatant was subsequently reduced, alkylated, and precipitated overnight in −20c freezer using trichloroacetic acid (15% total volume). Protein pellets were resuspended in 40 uL of loading buffer and briefly ran into a SDS-PAGE (approximately 5 mm., Invitrogen NuPAGE 4-12% Bis-Tris) for further purification, after which they were subjected to in-gel tryptic digestion using sequencing grade trypsin (Promega, V5113). After digestion, each sample was cleaned up using C18 columns and dried down. Peptides were then normalized using a nanodrop (ND-1000).

### Mass spectrometry

Peptide samples were diluted to 0.5 ug/uL. Subsequently, 1 uL was loaded onto an in-house laser-pulled 100 um ID nanospray column packed to ∼220mm with 3um 2A C18 beads (Reprosil). Peptides were separated by reversed-phase chromatography on a Dionex Ultimate 3000 HPLC. Buffer A of the mobile phase contained 0.1% formic acid (FA) in HPLC-grade water, while buffer B contained 0.1% FA in acetonitrile (ACN). An initial two-minute isocratic gradient flowing 3% B was followed by a linear increase up to 25% B for 115 minutes, then increased to 45% B over 15 minutes, and a final increase to 95% B over 15 minutes whereupon B was held for 6 minutes and returned back to baseline (2 min) and held for 10 minutes, for a total of 183 minutes. The HPLC flow rate was 0.400 uL/minute. Samples were run on either a Thermo Fusion Lumos (large study) or Thermo Orbitrap Elite (pilot comparisons) mass spectrometers that collected MS data in positive ion mode within the 400-1500 m/z range.

For TMT labeled samples, a top-speed MS3 method was employed on the Fusion Lumos with an initial Orbitrap scan resolution of 120,000. This was followed by high-energy collision-induced dissociation and analysis in the orbitrap using Top Speed’ dynamic identification with dynamic exclusion enabled (repeat count of 1, exclusion duration of 90 s). The automatic gain control for FT full MS was set to 4e5 and for ITMSn was set to 1e4. ITCID was used at MS2 method and the MS3 AGC was set to 1e5. The mass spectrometry proteomics data have been deposited to the ProteomeXchange Consortium via the PRIDE partner repository with the dataset identifier PXD017450 using ID: and password: kNvCO5eX.

### Peptide/Protein searches

#### Proteome Discoverer

The resulting mass spectra raw files were first searched using Proteome Discoverer 2.2. using the built-in SEQUEST search algorithm. Built-in TMT batch correction was enabled for all samples. Three FASTA databases were employed: Uniprot Swiss-Prot Homo sapiens (taxon ID 10090, downloaded January 2017), the Human Microbiome Project database (FASTA file downloaded from https://www.hmpdacc.org/hmp/HMRGD/ on January 2017), and a database containing common sample handling contaminants. Target-decoy searching at both the peptide and protein level was employed with a strict FDR cutoff of 0.05 using the Percolator algorithm built into Proteome Discoverer 2.2. Enzyme specificity was set to full-tryptic with static peptide modifications set to carmbamidomethylation (+57.0214 Da) and when appropriate, TMT (+229.1629 Da). Dynamic modifications were set to oxidation (+15.995 Da) and n-terminal protein acetylation (+42.011 Da). Only high-confidence proteins (q-val < 0.01) were used for analysis.

### Statistical analyses

Statistics were calculated using R with statistics packages (FactoMinerR 1.36, factoextra 1.0.5, ggplot2 2.2.1, Hmisc 4.0-3, psych 1.7.8, Mfuzz 2.34.0, ggpubr 0.1.5, RColorBrewer 1.1-2, UpSetR, 1.3.3, limma 3.30.13, venneuler 1.1-0), and Qlucore Omics Explorer 3.3. Protein abundance was normalized as a percentage of summed reporter intensity for all quantified proteins in a given sample (protein intensity / total sample intensity). Each TMT-11 run was filtered for. Where necessary for meeting statistical assumptions, abundances were log2 transformed. The appropriate multiple hypothesis tests (one-way ANOVA) were applied to abundance comparison data using Qlucore Omics Explorer or custom R scripts. Correlational p-values were corrected using false discovery rate (FDR) setting and the R package psych 1.7.8. Protein abundance heat maps were generated with Qlucore Omics Explorer 3.3 or R’s built-in heatmap function. FDRs and fold changes (where appropriate) were generated using Qlucore’s built-in FDR estimator, and the values are reported in supplementary tables.

## Author Contributions

SHT-Pro pipeline was designed by C.G.G. Proteomic sample preparation and mass spectrometry was conducted by C.G.G., H.C.W, and M.T. Samples were obtained by H.C.W., M.T., and J.L.S. Statistical analysis was done by C.G.G. and H.C.W. LOOCV Random Forest was generated by H.C.W. Manuscript writing and figure generation was done by C.G.G. and H.C.W., and edited by C.G.G., H.C.W., M.T., J.L.S., and J.E.E.

## Acknowledgements

The authors thank members of the Elias and Sonnenburg labs for their valuable input during the experimental and manuscript writing phases. The authors acknowledge the following sources of funding: C.G.G.: HHMI Gilliam Fellowship, Stanford Graduate Fellowship in Science and Engineering. C.G.G. and H.C.W.: NSF Graduate Fellowship (DGE – 114747). J.E.E.: Precision Health and Integrated Diagnostic (PHIND) Center at Stanford, and the Chan Zuckerberg Biohub. J.L.S.: NIH grants DK085025 and AT00989203. JLS is a Chan Zuckerberg Biohub Investigator.

## Conflicts of interest

The authors report no conflicts of interest

